# Truncated suPAR simultaneously causes kidney disease and autoimmune diabetes mellitus

**DOI:** 10.1101/2022.04.26.489589

**Authors:** Ke Zhu, Kamalika Mukherjee, Changli Wei, Salim S. Hayek, Agnieszka Collins, Changkyu Gu, Kristin Corapi, Mehmet M. Altintas, Yong Wang, Sushrut S. Waikar, Antonio C. Bianco, Jochen Reiser, Sanja Sever

## Abstract

Soluble urokinase-type plasminogen activator receptor (suPAR) is a risk factor for kidney diseases. Here we report the presence of C-terminal suPAR fragment, D2D3, in patients with diabetic nephropathy. D2D3-positive human sera inhibited glucose-stimulated insulin release in human islets and were associated with patients requiring insulin therapy. D2D3 transgenic mice presented kidney disease and diabetes marked by decreased levels of insulin and C-peptide, impaired glucose-stimulated insulin secretion, decreased pancreatic β-cell mass, and high fasting glucose. D2D3 fragment dysregulated glucose-induced cytoskeletal dynamics, impaired maturation and trafficking of insulin granules, and inhibited bioenergetics of β-cells in culture. An anti-uPAR antibody restored β-cell function in D2D3 transgenic mice. We show that the D2D3 fragment injures the kidney and pancreas, offering a unique dual therapeutic approach for kidney diseases and insulin-dependent diabetes.

**Summary:** Proteolytic suPAR fragment, D2D3, simultaneously injures two organs, the kidney and pancreas, thus causing a dual organ disease.

## Text

Chronic kidney diseases (CKDs) affect hundreds of millions of people worldwide. Three common causes of CKD are diabetes mellitus, hypertension, and glomerulonephritis (*1*). Diabetic nephropathy (DN) occurs in approximately 40% of diabetic patients and is a major cause of end-stage renal disease (ESRD) worldwide (*2*). The pathogenesis of DN encompasses diverse molecular mechanisms that include genetic, metabolic, and hemodynamic factors such as glomerular hypertrophy and hypertension (*2*). Histological and genetic data strongly implicates podocyte dysfunction in the development of DN (*3*).

Elevated levels of soluble urokinase-type plasminogen activator receptor (suPAR) have been associated with DN in patients with Type 1 diabetes (T1D) (*4*) and Type 2 diabetes (T2D) (*5–7*). suPAR is generated by proteolytic shedding of the membrane-bound uPAR from the surface of various cells of the innate immune system such as macrophages, immature myeloid cells, and neutrophils (*8, 9*). We and others have shown that uPAR/suPAR causes CKD by activating α_v_β_3_ integrin on glomerular podocytes, leading to foot process (FP) effacement and proteinuria (*10–14*).

uPAR/suPAR consists of three homologous domains (*15*): D1, D2, and D3. uPAR proteolysis generates two additional circulating forms (*16*): an N-terminal D1 fragment and a C-terminal D2D3 fragment, both of which are implicated in cancer biology (*15*). D2D3 fragment induces chemotaxis of cancer cells in part by activating α_v_β_3_ integrin (*15*); (*17*). Given that increased expression of β_3_ integrin has been associated with DN (*18*), we first sought to determine whether patients with DN have circulating D2D3 fragments (Fig. 1A).

**Figure 1.**
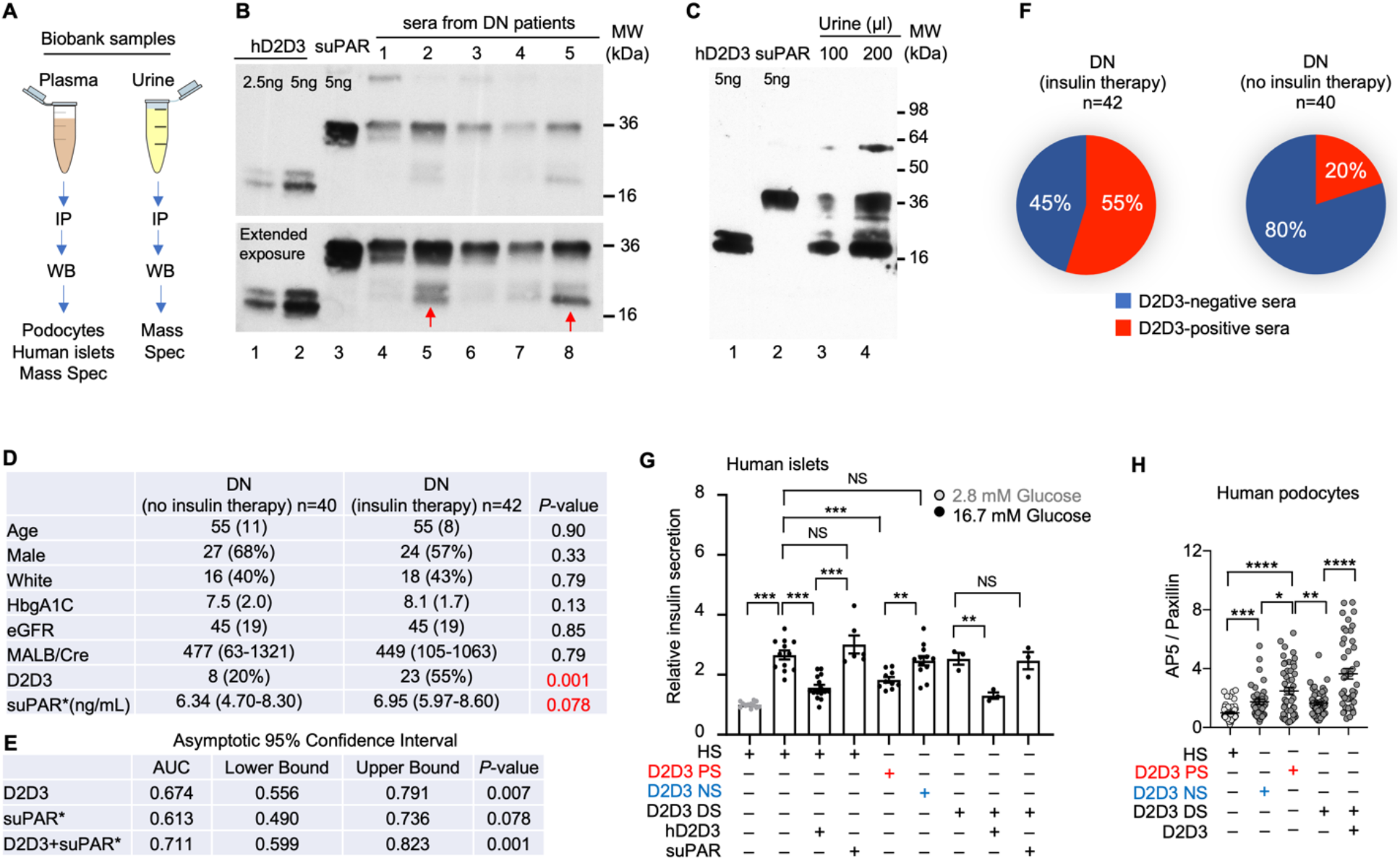
D2D3 fragment discriminates between DN patients on insulin therapy and those that are not. (**A**) Schematic of human sample analyses. (**B** and **C**) suPAR and D2D3 fragment detection by IP-WB in DN patient sera (**B**) or urine (**C**). Controls: recombinant human suPAR or chymotrypsin digested-suPAR representing D2D3 fragment. (**D**) Study patients’ characteristics. (**E**) Comparison (area under the curve, (AUC) values) between patients who were or were not on insulin therapy. (**F**) Pie chart showing the distribution (%) of D2D3-positive and D2D3-negative sera in DN patients who were or were not on insulin therapy. (**G**) Scatter dot plots representing levels of GSIS from human islets in the presence of human sera (n=3 experiments). Healthy donor islets were incubated in 10% healthy sera (HS), D2D3-positive sera (D2D3 PS, pooled from 8 patients), D2D3-negative sera (D2D3 NS, pooled from 6 patients), or D2D3 PS that were immunodepleted using an anti-uPAR antibody (R4) (D2D3 DS). When indicated, 10 ng/ml of recombinant hD2D3 or suPAR were added. (**H**) Scatter dot plots showing activation of β_3_ integrin on human podocytes (cells >35) treated with sera as described in (**G**), excluding hD2D3 concentration of 2.5 ng/ml. Immunofluorescence analysis was performed using an anti-paxillin antibody (focal adhesions), and an AP5 antibody (activated β_3_ integrin). **G** and **H**, error bar, mean ± SEM (\**P*≤0.05, \*\**P*≤0.01, \*\*\**P*≤0.001, \*\*\*\**P*≤0.0001, unpaired *t*-test). ns, not significant.

The presence of D2D3 was examined by modifying immunoprecipitation (IP)-coupled to Western blot (WB) analysis originally developed for cancer patients (fig. S1A) (*16, 19*). As expected, high levels of suPAR were detected in DN patient sera. Importantly, we identified proteins that were similar in molecular weight to proteolytically generated human D2D3 (hD2D3) fragments in sera of a subset of DN patients (Fig. 1B). We also detected hD2D3-like protein in the urine of DN patients (Fig. 1C), as seen previously in the urine of cancer patients (*20*). Of note, we and others did not detect hD2D3 in sera or urine from healthy individuals (fig. S1A) (*16, 19*). In order to demonstrate that our IP-WB analysis was indeed detecting hD2D3 fragment, we performed mass spectrometry (MS) analysis on proteins IPed from serum samples pooled from fourteen different DN patients (fig. S1B). MS identified a suPAR domain 2-specific peptide in the protein band corresponding to the D2D3 fragment’s molecular weight (figs. S1B, D and E). Additionally, WB followed by MS detected five suPAR-specific fragments in urine pooled from four DN patients (figs. S1C, D and F). Together, these data provide evidence for the presence of hD2D3 in a subset of DN patients.

DN is a consequence of T1D or T2D. In addition, a growing number of adults are diagnosed with late-onset insulin-dependent diabetes, often referred to as Latent Autoimmune Diabetes in Adults (LADA) (*21*). As T1D and LADA in adults are highly heterogeneous and are often misdiagnosed as T2D, we focused on DN patients who were either receiving or not receiving insulin therapy and investigated whether hD2D3 was preferentially present in one of the two patient populations. We analyzed two groups of patients matched in age, sex, glycated hemoglobin A1c (HgbA1c) levels, estimated glomerular filtration rate (eGFR), and urine albumin to creatinine ratio (MALB/Cre) (Fig. 1D). Circulating hD2D3 was present in 55% of DN patients requiring insulin therapy in contrast to only 20% of DN patients not requiring insulin therapy (Fig. 1F). In a logistic regression model adjusting for HgbA1c and suPAR, the presence of hD2D3 was associated with a 4.28-fold (95%CI[1.54-11.88]) increase in the odds of the patient being on insulin. The presence of the hD2D3 fragment in serum was a good discriminator between DN patients on insulin and those not on insulin (area under the curve (AUC) = 0.674) (Fig. 1E). We then elected to measure suPAR levels of the same patient population to investigate if the levels of D2D3 and full-length suPAR together can better distinguish between the insulin-dependent and independent DN patients. We first expressed and purified recombinant hD2D3 (fig. S2, A and B), and used it as a control to determine whether currently available suPAR ELISAs can detect hD2D3 fragment. Both commercially available ELISAs detected the recombinant hD2D3 fragment in addition to full-length suPAR (fig. S2C), suggesting that these assays report, when present, a combined level of the fragment and the full-length protein in sera. Using the same logistic regression model, we found that suPAR levels measured by ELISA together with the D2D3 fragment identified using IP-WB had better discrimination ability than either one of them individually (Fig. 1E). Together, these data suggest that circulating hD2D3 could be a previously unrecognized molecular link between kidney disease and insulin-dependent diabetes.

It is well established that the loss of β-cell function and decreased **g**lucose-**s**timulated **i**nsulin **s**ecretion (GSIS) underlie the pathogenesis of insulin-dependent diabetes. Thus, we next examined whether the presence of hD2D3 in sera would inhibit the GSIS of human islets. To be able to perform multiple assays using an identical set of sera, we pooled sera from 8 DN patients on insulin therapy who contained hD2D3 fragment (D2D3-positive sera) and 6 sera from DN patients who did not require insulin therapy (D2D3-negative sera). Compared to sera from healthy donors (HS), D2D3-positive sera impaired GSIS in human islets (Fig. 1G). This phenotype was reversed after hD2D3 and suPAR were immunodepleted using anti-uPAR antibody (D2D3 DS). Re-addition of recombinant hD2D3 but not suPAR resulted in impaired GSIS, convincingly demonstrating a direct role of hD2D3 in blocking insulin release by β-cells.

We have shown that suPAR induces glomerular disease by activating α_v_β_3_ integrin on podocytes (*22*). We next used immunofluorescence (IF) microscopy to test whether the presence of hD2D3 would mediate the ability of sera to activate β_3_ integrin on human podocytes. DN sera activated β_3_ integrin regardless of the presence of hD2D3 when compared to the HS, possibly due to higher levels of suPAR (Fig. 1H). Immunodepleting suPAR and hD2D3 abolished the activating phenotype, which was re-established upon the addition of the hD2D3 fragment (Fig. 1H). Similarly, the addition of hD2D3 converted non-activating HS into the activating serum in a concentration-dependent manner and was sufficient to induce β_3_ integrin activation in serum-free media (SFM) (Fig. S2D-F). These observations implicate D2D3 in podocyte injury that underlies DN and suggest that D2D3 may directly injure two organ-specific cells, glomerular podocytes and pancreatic β-cells.

To test the hypothesis that D2D3 can injure both the glomerulus and the pancreas, we generated a D2D3 transgenic mouse (D2D3-Tg), in which the expression of Myc-tagged mouse D2D3 (mD2D3, Fig. S2B) was driven by the AP2 promoter (Fig. 2A). D2D3-Tg animals were viable, fertile, and born at a normal Mendelian ratio. RT-qPCR detected elevated mD2D3-specific mRNA levels in fat tissues with no alteration for the D1-specific mRNA levels (Fig. 2B). Immunohistochemistry using an anti-Myc antibody detected the presence of Myc-tagged mD2D3 in the fat tissue (Fig. 2C). The mouse suPAR-specific ELISA detected an approximately 2-fold increase in the suPAR levels in the sera of transgenic animals (Fig. 2D). To check if mouse suPAR-specific ELISA detected both mD2D3 and suPAR in sera, we expressed and purified recombinant mouse Myc-tagged suPAR and mD2D3 as controls (Fig. S3A-C). Indeed, both suPAR and mD2D3 were detected by ELISA, albeit with lower sensitivity (fig. S3D). To identify and quantify the level of mD2D3 in circulation, we performed IP-WB and detected mD2D3 using an anti-Myc antibody. mD2D3 was detected at a concentration of 12±8 ng/ml only in the sera of D2D3-Tg, but not in control animals (Fig. 2E).

**Figure 2.**
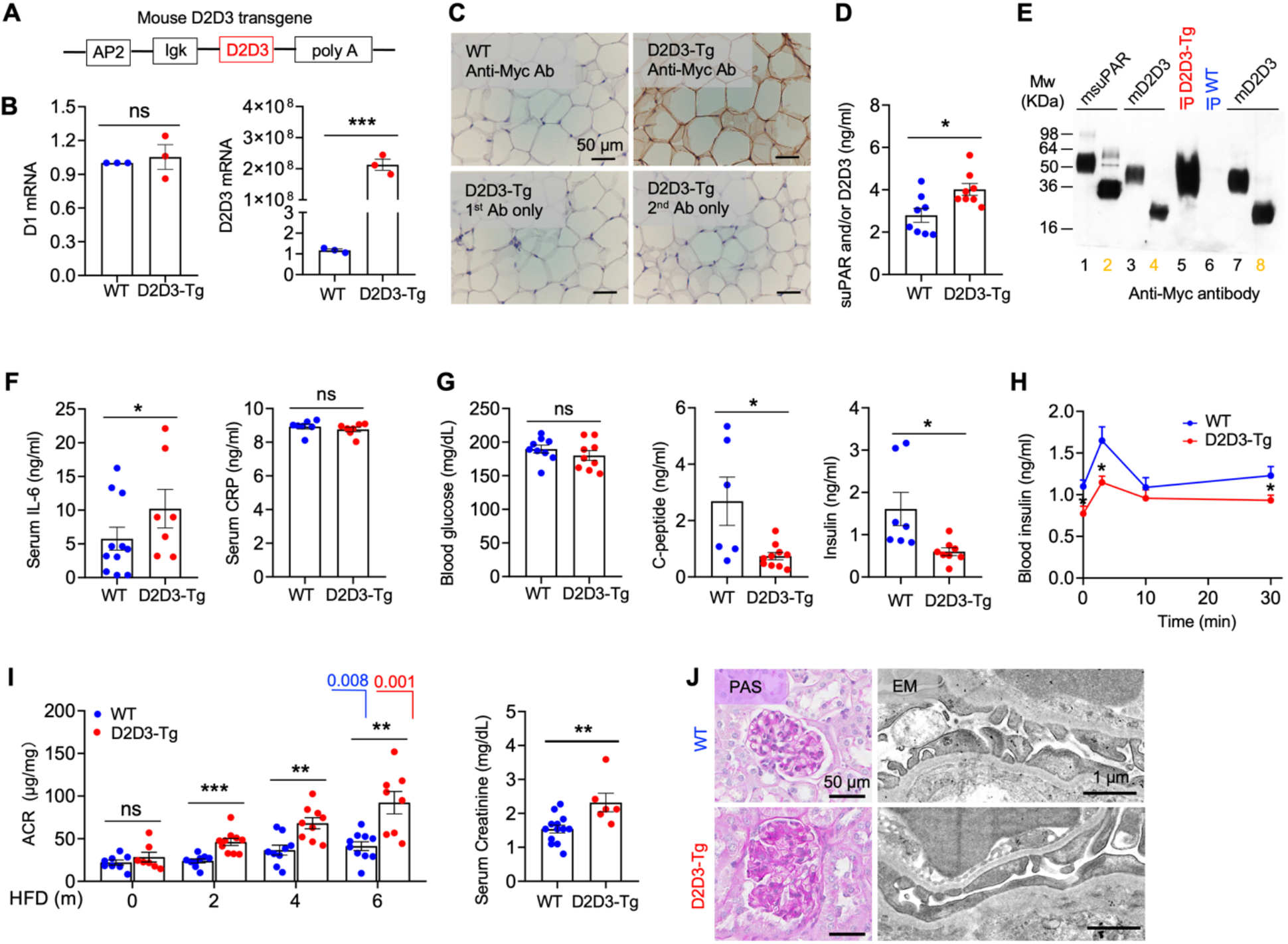
D2D3-Tg mice present glomerular injury and insulin insufficiency. All animals were fed a high-fat diet (HFD) and measurements were performed on 6 months old animals unless stated differently. (**A**) Schematics of D2D3-Tg construction. (**B**) qPCR analysis of *D2D3* and *D1* in fat tissues normalized to *Gapdh* (*n* = 3 biological replicates). (**C**) D2D3 detection in fat tissue of D2D3-Tg mice using anti-Myc antibody (D2D3 is Myc tagged). Controls: tissues stained with primary or secondary antibody only. (**D**) Mouse serum suPAR and D2D3 fragment levels were determined using uPAR-specific ELISA (*n* = 8). (**E**) Detection of Myc-tagged D2D3 in mouse sera. Sera were IPed with anti-uPAR antibody and detected by WB using anti-Myc antibody. Control: recombinant Myc-tagged mD2D3 (Lane 2, 6; +/- N-glycanase treated). (**F**) Serum IL-6 (WT, *n* = 11; D2D3-Tg, *n* = 7) and CPR (*n* = 7, each group) levels. (**G**) Fasting blood glucose (*n* = 9, each group), C-peptide (*n* = 6 for WT; *n* = 10 for D2D3-Tg), and insulin (*n* = 7 for WT; *n* = 8 for D2D3-Tg) levels. (**H**) *In vivo* GSIS. Overnight fasted animals were intraperitoneally injected with glucose (2 g/kg body weight) and their blood insulin levels were measured by ELISA (*n* = 6-8 for each group). (**I**) Scatter dot plots presenting albumin creatinine ratio (ACR) or serum creatinine (*n* = 7-10 for each group) levels. (**J**) PAS stained kidney (left) and TEM of foot processes (right). (**C, J**). For all results, error bar, mean ± SEM, (\**P* < 0.05; \*\**P* < 0.01; \*\*\**P* < 0.001. unpaired *t*-test). ns, not statistically significant.

In line with the presence of hD2D3 fragment in sera from DN patients, D2D3-Tg animals exhibited higher levels of IL-6 (Fig. 2F), a pro-inflammatory cytokine often increased in patients with diabetes (*23*). Although levels of serum CRP or blood glucose were not significantly increased (Fig. 2F), D2D3-Tg animals exhibited lower levels of insulin and C-peptide suggesting impaired pancreatic β-cell function (Fig. 2G). As observed using D2D3-positive human sera and human islets, D2D3-Tg mice exhibited impaired GSIS (Fig. 2H). This phenotype was unique for mD2D3, as transgenic mice overexpressing full-length mouse suPAR (suPAR-Tg) (*24*) did not exhibit decreased levels of insulin (fig. S4A). These data further confirm the role of the D2D3 fragment in inhibiting β-cell function.

Since we observed hD2D3 dependent activation of α_v_β_3_ integrin on human podocytes, we investigated the renal health of D2D3-Tg mice. Indeed, D2D3-Tg animals developed glomerular injury similar to a pattern seen in DN: progressive microalbuminuria accompanied by FP effacement, elevated serum creatinine, glomerular hypertrophy, and mesangial expansion (Fig. 2, I and J). Together with our data using human sera, these studies provide compelling evidence for the direct role of proteolytic D2D3 fragment in inducing glomerular injury by activating α_v_β_3_ on podocytes, and insulin insufficiency by impairing the function of β-cells.

The phenotypic analysis in Figure 2 was performed on 6 months-old mice fed with a high-fat diet (HFD) that enabled the expression of higher levels of mD2D3. As HFD can induce chronic low-grade systemic inflammation (*25*), we next examined the effects of mD2D3 on animal physiology when animals were fed a regular diet. As expected, there was a significant difference in the bodyweight of animals on different diets (fig. S4B). Even on a regular diet, D2D3-Tg mice expressed mD2D3 fragment (fig. S4, C and D), with suPAR/D2D3 levels increasing as animals aged and gained weight (fig. S4, E and F). Interestingly, D2D3-Tg mice on a regular diet did not develop significant kidney injury phenotypes (fig. S4G) in contrast to the DN type of glomerular injury on HFD. This suggests that either kidney injury requires the presence of higher levels of mD2D3 fragment in circulation, or that mD2D3 fragment needs to synergize with HFD to induce kidney injury.

Although diet affected the D2D3-mediated kidney phenotype, it had no influence on the mD2D3-induced pancreatic phenotypes. D2D3-Tg mice fed with a regular diet exhibited decreased levels of C-peptide and blood insulin, as well as compromised *in vivo* GSIS even at an early age of 2 months (Fig. 3, A and B). Islets isolated from the D2D3-Tg mice exhibited impaired *ex-vivo* GSIS (Fig. 3C), demonstrating that the observed phenotypes were inherent to the function of the pancreas. Concomitant with decreased insulin secretion, D2D3-Tg mice exhibited impaired glucose handling ability as evidenced glucose tolerance test (GTT) (Fig. 3D). Finally, the fasting blood glucose levels increased by 12 months of age (Fig. 3E). Together, these data show that D2D3-Tg mice develop insulin-dependent diabetes.

**Figure 3.**
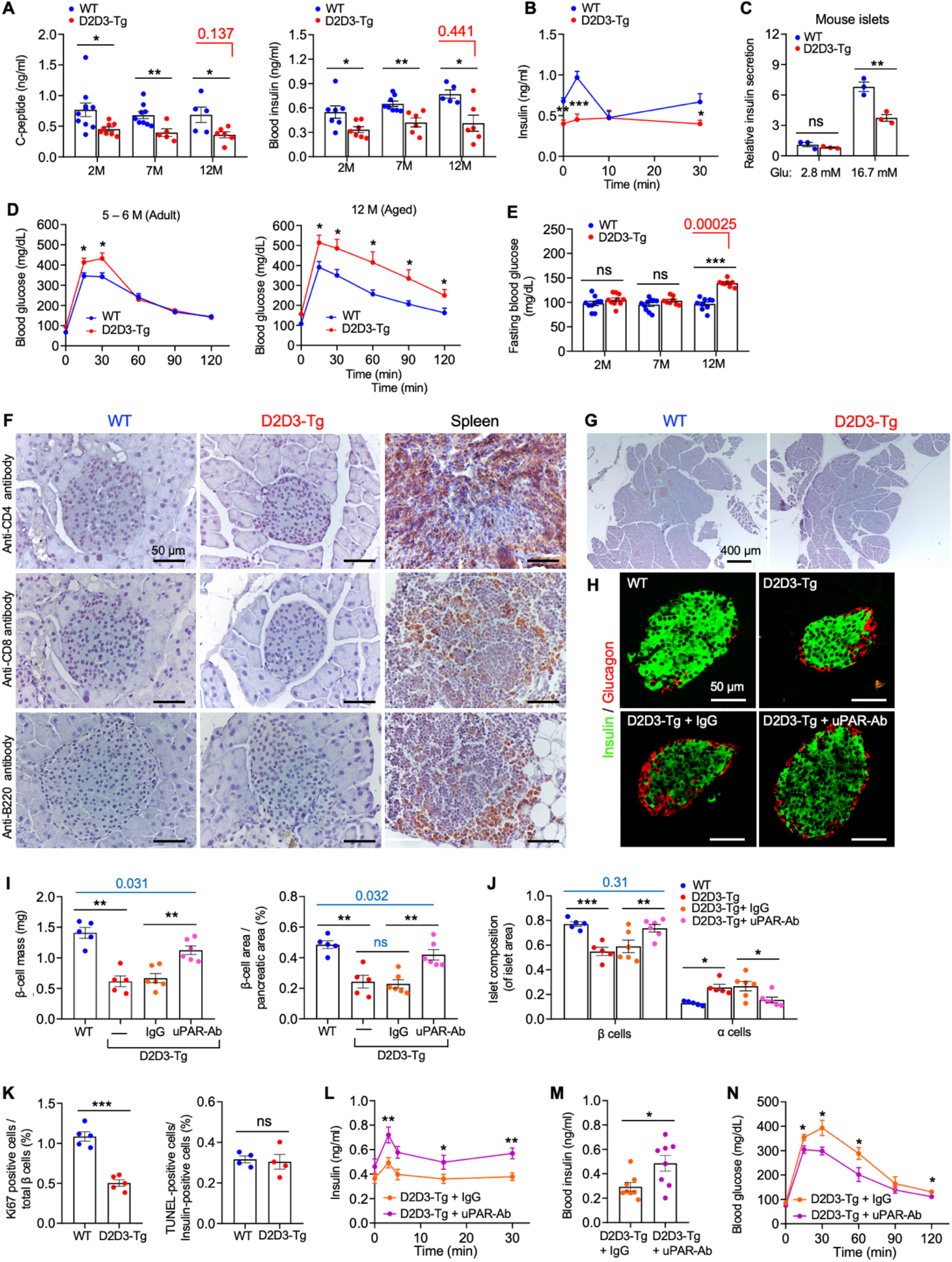
D2D3-Tg mice present insulin-dependent diabetes due to impaired pancreas function and β-cell mass. All animals were fed a regular diet. (**A**) Scatter dot plots representing fasting C-peptide and blood insulin levels (*n* = 5-10 mice in each group) (**B**) *In vivo* GSIS. Overnight fasted animals were intraperitoneally injected with glucose (2 g/kg body weight) and their blood insulin levels were measured by ELISA (WT, *n* = 5; D2D3-Tg, *n* = 8). (**C**) *In vitro* GSIS. Islets from 2-month-old animals (*n* = 3) were subjected to *in vitro* GSIS. (**D**) Glucose tolerance test (GTT). Adult (5∼6 months) and aged (12 months) mice were fasted overnight before intraperitoneally injected with glucose (2 g/kg body weight). Blood glucose was measured at the indicated time points (*n* = 5-6 animals per each condition). (**E**) Scatter dot plots depicting fasting blood glucose levels (*n* = 8-10 mice per condition). (**F**) Representative immunohistochemistry of islets from 2-month-old mice stained with anti-CD4,anti-CD8, and anti-B220 antibodies. Spleen was used as a positive control for T and B cells staining (*n* = 4 mice per each condition). (**G**) Representative immunohistochemistry of pancreas from 2-month-old mice stained with anti-insulin antibody (*n* = 5 mice per each condition). (**H**) Representative immunohistochemistry of islets using anti-glucagon (α-cell) and anti-insulin (β-cell) antibodies (*n* = 5 mice per each condition). (**I**) Scatter dot plots representing β-cell mass and β-cell area/pancreatic area ratio. Data were generated using images shown in (**H**). When indicated, D2D3-Tg mice were treated with anti-uPAR-Ab or IgG isotype control (IgG) for four weeks beginning at 2 months of age. Those data were collected at 3-month-old animals. (*n* = 5 for WT and D2D3-Tg; *n* = 6 for D2D3-Tg treated with either IgG or uPAR-Ab). Between 6-8 sections from each tissue were analyzed. (**J**) Scatter dot plots representing the composition of the islets in animals treated as described in (**I**). Islet composition was determined by counting the total number of β-cells (green) and α-cells (red) and expressing them as percentages of total cells counted within the single islet. (**K**) Scatter dot plots depicting levels of apoptosis determined by TUNEL staining, or cell proliferation determined by Ki67-positive staining of mice islets (*n* = 5 mice for each condition). Each dot represents an average of 1000-1350 insulin-positive cells counted per animal. (**L**) *In vivo* GSIS using two months old D2D3-Tg treated with either IgG or anti-uPAR-Ab twice within one week. All animals were male (*n* = 7-10 animals for each condition). (**M** and **N**) D2D3-Tg mice were treated with either anti-uPAR-Ab or IgG beginning at 2 months of age for four weeks. Experiments were performed at 3 months of age. All animals were male. Scatter dot plots representing blood insulin levels are shown in (**M**) (*n* = 8 mice for each condition). The glucose tolerance test is shown in (**N**) (*n* = 6 mice for each condition). For all results, error bar, mean ± SEM, (\**P* < 0.05; \*\**P* < 0.01; \*\*\**P* < 0.001. unpaired *t*-test). ns, not statistically significant. In addition, when appropriate *P* numbers are shown in red or blue.

One of the characteristics of insulin-dependent diabetes is the reduction of pancreatic β-cells in part due to infiltration of autoreactive CD4 (+) and CD8 (+) T-cells (*26*). Interestingly, D2D3-Tg animals did not exhibit major infiltration of CD4 (+), CD8 (+) T-cells, or B220 (+) B-cells either at 2-months of age (Fig. 3F) or at 12-months of age (fig. S4H), indicating the involvement of a different mechanism that governs D2D3-mediated injury. Thus, we further examined the islet architecture and cell distribution by staining the pancreas for insulin (β-cell marker) and glucagon (α-cell marker) (Figs. 3, G and H). Although, the D2D3-Tg animals did not exhibit decreased overall pancreatic weight (fig. S4I), surprisingly, we observed lower β-cell area and reduced β-cell mass (Fig. 3I). Additionally, a higher percentage of the α-cell population was observed, which altered the overall composition of the islets (Fig. 3J). Of note, these phenotypes were observed only in the post-natal animals, as neonatal wild type and D2D3-Tg mice exhibited similar distribution of pancreatic β-cells and α-cells (fig. S5A-C). Given that the reduced β-cell mass is only observed in post-neonatal D2D3-Tg animals, we next examined the proliferative capability of β-cells using Ki67 marker (*27*), as well as the level of apoptosis using TUNEL staining of the mouse islets (*27*). While we found no difference in the level of apoptotic cells in the islets of wild type and D2D3-Tg mice, β-cell proliferation was impaired in the D2D3-Tg mice (Fig. 3K). Together, these data demonstrate that the presence of D2D3 in circulation impairs β-cell proliferation, which leads to lower β-cell mass in postnatal D2D3-Tg mice in comparison to its wild-type counterpart.

To further affirm that the observed pancreatic injury phenotypes are a direct effect of D2D3, we tested whether neutralizing the circulating D2D3 by anti-uPAR antibody(uPAR-Ab) can ameliorate the injury phenotypes. We first treated 2 months old age-matched male D2D3-Tg mice twice within one week with either mouse anti-uPAR or IgG (control) antibody. Treatment with uPAR-Ab significantly restored the islet function in D2D3-Tg mice, demonstrated by improved *in vivo* GSIS (Fig. 3L). We continued to treat mice twice a week for the following four weeks and observed that uPAR-Ab treated animals exhibited an increase in fasting blood insulin levels, and improved glycemic excursion during glucose tolerance test in uPAR-Ab treated D2D3-Tg mice (Fig. 3, M and N). The observed positive effect of uPAR-Ab on the function of the pancreas was due to improved β-cell area and mass as well as overall islet composition (Fig. 3, I and J). Together, these data identify a novel mechanism that underlies insulin-dependent diabetes mellitus in which circulating D2D3 fragment directly injures β-cells of the pancreas, and suggest that this pathogenic mechanism can be ameliorated by blocking D2D3 fragment using specific anti-uPAR-Ab.

We used MIN6 cells, a cell line derived from a mouse insulinoma with characteristics of pancreatic β-cells (*28*), to further elucidate the molecular mechanism by which D2D3 affects β-cell physiology. The addition of recombinant mD2D3 impaired GSIS of MIN6 cells in a dose- and time-dependent manner (fig. S5D), and this effect was inhibited by the addition of anti-uPAR-Ab (Fig. 4A). This effect was D2D3-specific since the addition of full-length suPAR did not inhibit GSIS (Fig. S5D). Identical trends were observed in isolated mouse islets, as well as human islets (Fig. 4B, C). Together, these data demonstrate that D2D3 directly impairs the ability of β-cells to release insulin upon glucose stimulation and establishes MIN6 cells as a relevant cell line for studying the effect of D2D3 on insulin secretion.

**Fig 4.**
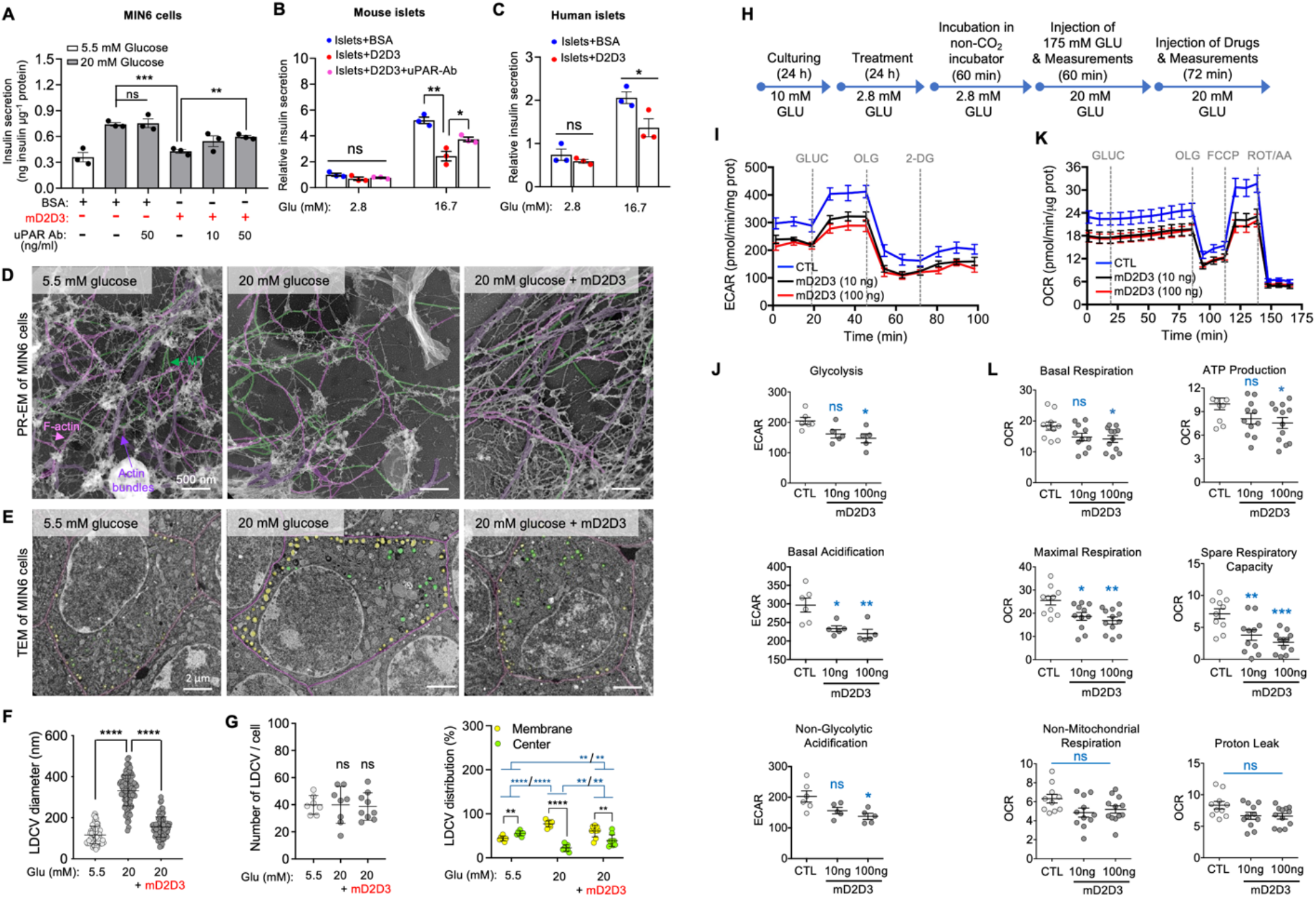
D2D3 fragment impairs multiple aspects of β-cell physiology. (**A**) Graphs depicting GSIS by MIN-6 cells were incubated in the presence or absence of BSA, mD2D3 (100 ng/ml), and anti-uPAR Ab (*n* = 3). (B) Graphs depicting GSIS by mouse islets were incubated in the presence or absence of BSA, mD2D3 (100 ng/ml), and anti-uPAR Ab (*n* = 3). (C) Graphs depicting GSIS by human islets were incubated in the presence or absence of BSA, hD2D3 (100 ng/ml) (*n* = 3). (**D**) PR-EM micrographs showing the cytoskeleton of MIN6 cells. MIN6 cells were grown as described in (**A**) with exception of increasing glucose levels to 20 nM. When indicated, an mD2D3 fragment (100 ng/ml) was added for 24 hours. Actin bundles (pink), hyper bundles (purple), and microtubules (MT, green). (**E**) TEM micrographs of MIN-6 cells were treated as described in (D). Large dense-core vesicles (LDCVs) localized near (yellow) the plasma membrane (pink) and those localized more than 500 nm away (green) from the plasma membrane. (**F**and **G**) Graphs represent average LDCV diameter (**F**), number of LDCV per cell, and size based on location (**G**) in cells shown in (**E**). In (F), more than 75 vesicles were counted in ≥10 cells per treatment. Of note, the overall number of LDCV per cell regardless of the treatment was similar and ranged between 17-57. Error bar, mean ± SD (\*\**P* < 0.01; \*\*\**P* < 0.001, unpaired *t-*test). (**H**) Schematic for the experimental setup for glycolysis and the oxygen consumption rate (OCR) measurement in MIN-6 cells. (**I-J**) Extracellular acidification rates (ECAR) were measured in untreated (CTL) and D2D3-treated cells using the Seahorse XFe24 analyzer. When indicated, cells were treated with hD2D3. (**K-L**) OCR curves and measurements of the bioenergetic parameters regarding mitochondrial respiration of the treated cells as explained in (**H**). Each OCR value was normalized to cell number and presented as pmol/min/100,000 cells (three experiments). For **J** and L Error bars mean ± S.E.M. (*P<0.05, \*\**P*≤0.01, \*\*\**P*≤0.001, one-way ANOVA). ns, not statistically significant.

To determine which steps in insulin release were inhibited by mD2D3, we next examined the effect of D2D3 had on the cytoskeleton using platinum replica electron microscopy (PR-EM) (fig. S6). It is well documented that high glucose leads to microtubules (MT) polymerization, which aids the transport of secretory vesicles from the cell periphery to the plasma membrane, and simultaneous depolymerization of actin networks (*29*). Our limited analysis suggested that mD2D3 inhibited both processes: impaired MT polymerization and inhibited actin networks depolymerization (Fig. 4D). As dysregulated cytoskeletal dynamics is expected to have a profound effect on the trafficking and maturation of large dense-core vesicles (LDCVs) that transport insulin in β cells, we next visualized LDCVs using transmission electron microscopy (TEM) (fig. S7). As seen in β-cells (*30, 31*), glucose stimulation increased the diameter of LDCVs (vesicle maturation) as well as the number of membrane-associated LDCVs (vesicle trafficking) (Fig. 4E-G). Addition of mD2D3 impaired both of these processes demonstrating that the mD2D3 fragment impairs multiple steps of insulin secretion by dysregulating glucose-induced cytoskeleton dynamics.

As insulin secretion is also directly coupled to glycolysis and oxidative metabolism (*32, 33*), we examined whether mD2D3 affected glycolysis and mitochondrial respiration in MIN6 cells upon glucose stimulation. To this end, extracellular acidification (ECAR) and oxygen consumption rates (OCR) were measured in real-time in MIN6 cells under basal conditions and in response to sequential injections of glycolysis or mitochondrial inhibitors using a Seahorse XFe24 extracellular flux analyzer. To mimic glucose stimulation, cells were grown at low glucose (2.8 mM) for 24 hours in the presence of increasing concentrations of D2D3 prior to switching to high glucose (20 mM) and initiation of the experiment (Fig. 4H). The presence of mD2D3 decreased glycolysis, basal acidification, and non-glycolytic acidification (Fig. 4I and J). In addition, mD2D3 decreased several bioenergetic parameters including basal and maximal respiration, spare respiratory capacity, and ATP production (Fig. 4K and L). Inability to increase OCR in response to glucose is expected to impair insulin granule maturation and trafficking as well as the proliferative capacity of β-cells. Therefore, these data identify a multi-consequential novel molecular mechanism by which a circulating pathogenic factor, D2D3, alters glucose-dependent β-cell physiology, including glycolysis, OCR, insulin granule maturation and trafficking, cytoskeletal dynamics, insulin secretion, and ultimately impairs β-cell proliferation in the post-natal period.

Among the US population, 34.2 million people of all ages have diabetes. Although T2D is the most prevalent form of diabetes, a growing number of adults are diagnosed with either T1D or LADA. Originally it was suggested that the main difference between T1D and LADA is the age at which diseases present: T1D commonly occurs in children, whereas LADA is diagnosed in adulthood. Recent data suggest that the main difference is not the age but the speed by which these diseases progress, thus implicating distinct molecular mechanisms for each disease (*34, 35*). Despite these differences, a loss of pancreatic β-cells is a central feature of diabetes regardless of its etiology. Recently, signaling by the death receptor TMEM219 expressed on β-cells via its circulating ligand insulin-like growth factor binding protein 3 (IGFBP3) has been implicated in the loss of β-cells via induction of apoptosis (*36*). It was suggested that blocking IGFBP3/TMEM219 signaling pathway might present a novel therapeutic option for both T1D and T2D. Our study identifies a novel molecular mechanism of diabetes in which a different circulating protein, the D2D3 fragment of suPAR, causes injury to the β-cells by impairing insulin secretion and β-cell proliferation in the post-natal period via inhibition of glucose-dependent cell metabolism. Furthermore, in contrast to β-cell-specific mechanisms of injury (*36, 37*), our study shows that the D2D3 fragment is capable of injuring two organs simultaneously. Treatment with uPAR-Ab attenuated D2D3-driven pancreatic injury in *in vivo* and *ex vivo* mouse models, and immunodepletion of hD2D3 from DN patient sera abated pathogenic phenotypes in human podocytes and islets. We thus envision the possibility of using D2D3-specific antibodies as a unique dual therapeutic option for CKD and insulin-dependent diabetes.

D2D3 fragment is generated by the proteolysis of uPAR, which is an effector of the innate immune system as it is expressed by macrophages, neutrophils, and immature myeloid cells (*14*). Therefore, our study identifies a mechanism through which chronic inflammation leads to autoimmune diabetes mellitus. Given the central role of suPAR as a major risk factor in the COVID-19 associated kidney injury (*38, 39*), as well as adverse outcomes in patients with diabetes mellitus (*40*), our study warrants caution for the increased appearance of innate autoimmune diabetes in the post-COVID period globally.

## Acknowledgments

We would like to thank Elizabeth Ankers for her help in collecting samples at Renal Associates, Boston. We thank Bradley P. Pedro for technical assistance with immunocytochemistry and the quantification of podocyte phenotypes. We thank Decheng Ren, University of Chicago, for providing us with MIN6 cells. We acknowledge the support from the National Institutes of Health: R01 DK093773 and DK087985 to S.S., DK125858 and DK101350 to J.R., HL153384 and DK128012 to S.S.H. We acknowledge the help from the Electron Microscopy and the Taplin Biological Mass Spectrometry facilities at Harvard Medical School. We would like to thank Susan Bonner-Weir, Joslin Diabetes Center/Harvard Medical School, for her constructive comments on the manuscript.

## Author contributions

J.R. and S.S initiated, designed, and oversaw the research. S.S, K.Z. and K.M. assembled the figures and wrote the paper. K.Z. expressed and purified recombinant proteins, performed GSIS using islets and MIN6 cells, and all the animal experiments. K.M. set up IP-WB, analyzed human and mouse samples, generated samples for mass spectrometry, and carried out cell-based assays using human sera. C.W. oversaw all animal experiments and performed histological analysis of tissue samples. S.S.H. performed statistical analysis of human data sets. A.C. carried out EM experiments. C.G. performed cell culture experiments using recombinant proteins. K.C. collected human samples while she worked for Renal Associates at MGH. S.S.W. provided human samples from the MGB biobank. M.M.A. carried out Seahorse extracellular flux experiments. Y.W. provided human islets. A.B. discussed results and provided insights regarding human endocrinology. All authors discussed the results.

