## Supplemental material for "Truncated suPAR simultaneously causes kidney disease and autoimmune diabetes mellitus"

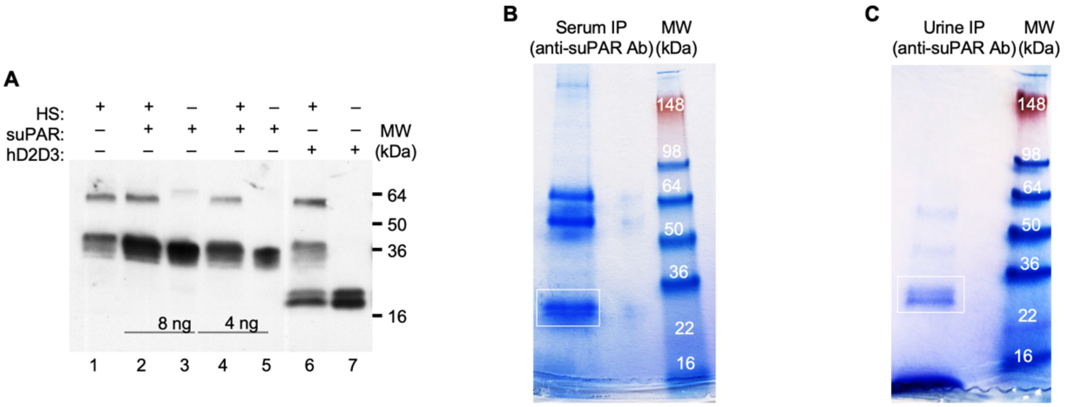


**D**

**Domain I** Fragment-**1**
MGHPPLLPLLLLLHTCVPASWGLRCMQCKTNGDCRVEECALGQDLCRTTIVR**LWEEGEELELVEK**SCTHSEKTNRTLSYRTGLKITSLTEVVCGLDLCNQGNSGRA

**Domain II** Fragment-**2** Fragment-**3/3**
VTYSRSRYLECISCGSSDMSCERGRHQSLQCR**SPEEQCLDVVTHWIQEGEEGRPK**DDRHLRGCGYLPGCPGSNGFHNNDTFHFLKCCNTTK**CNEGPILELENLPQ**

**Domain III** Fragments-**4/4*** Fragment-**5**
**NGR**QCYSCKGNSTHGCSSEETFLIDC**RGPMNQCLVATGTHEPK**NQSYMVRGCATASMCQHAHLGDAFSMNHIDVSCCTK**SGCNHPDLDVQYR**SGAAPQPGPAH

**GPI**
LSLTITLLMTARLWGGTLLWT

**E**

suPAR fragment IP-ed from sera

| ScanF | z | ↓XCorr | ΔCorr | # Ions | Reference | Redun | Peptide | Gene Symbol | Max | Fragment # |
|---|---|---|---|---|---|---|---|---|---|---|
| 18271 | 4 | 4.761 | 0.402 | 40/132 | sp\|Q03405\|UPAR_HUMAN | 7 | R.SPEEQCLDVVTHWIQEGEEGRPK.D | PLAUR | 38428.9 | 2 |

**F**

suPAR fragments IP-ed from urine

| ScanF | z | ↓XCorr | ΔCorr | # Ions | Reference | Redun | Peptide | Gene Symbol | Max | Fragment # |
|---|---|---|---|---|---|---|---|---|---|---|
| 12288 | 4 | 5.226 | 0.491 | 44/132 | sp\|Q03405\|UPAR_HUMAN | 7 | R.SPEEQCLDVVTHWIQEGEEGRPK.D | PLAUR | 74129.3 | 2 |
| 11068 | 2 | 4.563 | 0.326 | 21/24 | sp\|Q03405\|UPAR_HUMAN | 6 | R.LWEEGEELELVEK.S | PLAUR | 52916.7 | 1 |
| 4638 | 3 | 3.597 | 0.2 | 24/60 | sp\|Q03405\|UPAR_HUMAN | 0 | R.GPMNQCLVATGTHEPK.N | PLAUR | 25545.2 | 4 |
| 11488 | 2 | 3.448 | 0.398 | 16/32 | sp\|Q03405\|UPAR_HUMAN | 5 | K.CNEGPILELENLPQNGR.Q | PLAUR | 4710.2 | 3/3 |
| 2958 | 3 | 2.71 | 0.371 | 25/60 | sp\|Q03405\|UPAR_HUMAN | 0 | R.GPM*NQCLVATGTHEPK.N | PLAUR | 62054.6 | 4* |
| 4122 | 2 | 1.864 | 0.224 | 14/24 | sp\|Q03405\|UPAR_HUMAN | 1 | K.SGCNHPDLDVQYR.S | PLAUR | 3435.8 | 5 |
| 3038 | 2 | 1.473 | 0.011 | 30-Dec | sp\|Q03405\|UPAR_HUMAN | 0 | R.GPM*NQCLVATGTHEPK.N | PLAUR | 7125.7 | 4* |

**S1. D2D3 fragment is present in a subset of DN patients**.

(**A**) Sera from healthy individuals do not contain D2D3 fragment. Where indicated, recombinant human suPAR (lanes 2- 5), or hD2D3 fragments generated by chymotrypsin digestion of full-length suPAR (lanes 6 and 7) were added to healthy human sera (HS) (lanes 2, 4 and 6). suPAR proteins were immunoprecipitated (IPed) from serum samples using monoclonal anti-uPAR antibody (R4), and deglycosylated using N-glycanase before performing western blot (WB) analysis using a polyclonal anti-uPAR antibody. Note that HS contains full length suPAR (lane 1) but no D2D3 fragment. IP efficiency observed in this IP-WB assay for full-length suPAR and hD2D3 were 74±2% and 55±8%, respectively.

(**B**) Coomassie gel showing proteins IPed from sera using anti-uPAR antibody (R4). Serum samples (10 ml) from 14 DN patients that also had a D2D3-like fragment in their sera were used in the experiment. The protein band indicated in the figure was cut out of the gel and analyzed using mass spectrometry.

(**C**) Coomassie gel showing proteins IPed from urine using anti-uPAR antibody (R4). Urine samples (18 ml) from 4 DN patients were used in the experiment. The protein band indicated in the figure was cut out of the gel and analyzed using mass spectrometry.

(**D**) Amino acid sequence of human uPAR protein isoform 1. Distinct domains are color-coded in blue (D1), yellow (D2), and green (D3). Gray color marks the linker between D1 and D2, and the GPI anchor. Bold letters demarcate suPAR-specific peptides detected by mass spectrometry.

(**E, F**) Tables present suPAR-specific fragments that were identified in sera (**E**) or urine (**F**) using mass spectrometry. Numbers for the fragments correspond to the sequences marked in (**C**). Asparagine (N) in the fragment-4 may not be fully deglycosylated. *Additional fragment generated within the fragment-4. Fragment-3 spans over D2 and D3 domain and thus is marked 3/3. Data demonstrate that suPAR protein was present in the serum and urine of DN subjects.

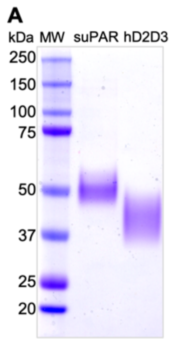


**B**

```
Human uPAR   15   TCVPASWGLRCMQCKTNGDCRVEECALGQDLCRTTIVRLWEEGEELELVEKSCTHSEKTN   74
                  TCVPAS GL+CMQC++N  C VEECALGQDLCRTT++R W++  ELE+V + C HSEKTN
Mice uPAR    16   TCVPASQGLQCMQCESNQSCLVEECALGQDLCRTTVLREWQDDRELEVVTRGCAHSEKTN   75
                                                                 hD2D3
Human uPAR   75   RTLSYRTGLKITSLTEVVCGLDLCNQGNSG-RAVTYSRSRYLECISCGSSDMSCERGRHQ   133
                  RT+SYR G  I SLTE VC  +LCN+   G R   + + RYLEC SC S D SCERGR Q
Mice uPAR    76   RTMSYRMGSMIISLTETVCATNLCNRPRPGARGRAFPQGRYLECASCTSLDQSCERGREQ   135
                                                                 mD2D3
Huma uPAR   134   SLQCRSPEEQCLDVVTHWIQEGEEGRPKDDRHLRGCGYLPGCPGSNGFHNNDTFHFLKCC   193
                  SLQCR P E C++VVT    +  E   KD+ + RGCG LPGCPG+ GFH+N TFHFLKCC
Mice uPAR   136   SLQCRYPTEHCIEVVT---LQSTERSLKDEDYTRGCGSLPGCPGTAGFHSNQTFHFLKCC   192

Human uPAR  194   NTTKCNEGPILELENLPQNGRQCYSCKGNSTHGCSSEETFLIDCRGPMNQCLVATGTHEP   253
                  N T CN GP+L+L++ P NG QCYSC+GN+T GCSSEE  LI+CRGPMNQCLVATG
Mice uPAR   193   NYTHCNGGPVLDLQSFPPNGFQCYSCEGNNTLGCSSEEASLINCRGPMNQCLVATGLDVL   252

Human uPAR  254   KNQSYMVRGCATASMCQHAHLGDAFSMN-HIDVSCCTKSGCNHPDLDVQ-Myc tag
                  N+SY VRGCATAS CQ +H+ D+F  + ++ VSCC  SGCN P
Mice uPAR   253   GNRSYTVRGCATASWCQGSHVADSFPTHLNVSVSCCHGSGCNSP------Myc tag
```

**C**

| Input | ELISA (R&D) | ELISA (Virogates) | IP-WB suPAR | IP-WB hD2D3 |
|---|---|---|---|---|
| hsuPAR (5 ng/ml) | 1.8 ng/ml | 3.2 ng/ml | NA | NA |
| hD2D3 (5 ng/ml) | 1.2 ng/ml | 5.1 ng/ml | NA | NA |
| HS serum | 1.7 ng/ml | 2.7 ng/ml | 5.8 ng/ml | NA |
| DN serum 1# | 2.2 ng/ml | 6.8 ng/ml | 16.3 ng/ml | 6.6 ng/ml* |
| DN serum 5# | 5.2 ng/ml | 12.3 ng/ml | 12.6 ng/ml | 12 ng/ml |

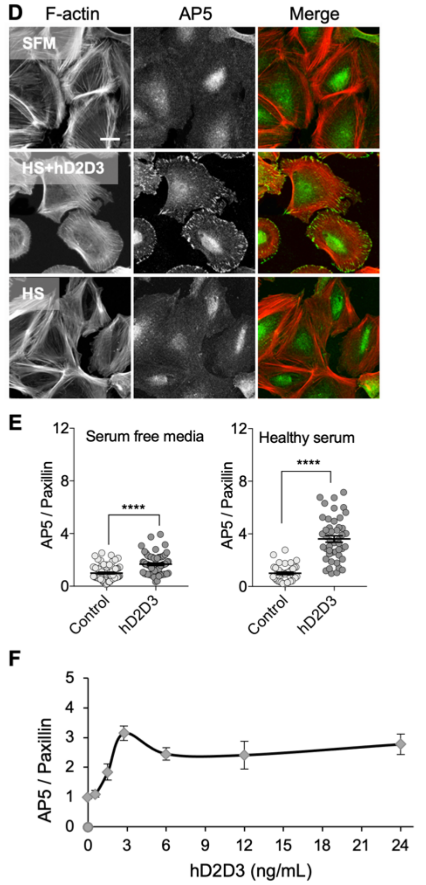


**Figure S2. Recombinant hD2D3 fragment activates β3 integrin on human podocytes.**

(**A**) Coomassie gel showing recombinant human suPAR and hD2D3 fragment that were expressed and purified from HEK-293 cells. Notice that protein purification resulted in highly pure proteins. As both proteins are glycosylated, they travel on a gel as wide bands.

(**B**) Alignment of amino acid sequences for human and mouse suPAR proteins. Recombinant suPARs and D2D3 fragments were Myc-tagged on their C-terminus. Tagging of the proteins removed GPI-anchor sequence, thus generating soluble proteins. The first amino acid of the mD2D3 expressed in D2D3-Tg mice was alanine 120 (labeled in orange).

(**C**) Two commercially available ELISAs detect both, the full length suPAR and the D2D3 fragment. Notice that Virogates assay is more sensitive than R&D assay, and that both ELISAs are less sensitive than the IP-WB analysis. #Data for DN sera were calculated from samples shown in Fig. 1B. Since majority of DN sera contained lower molecular weight proteins similar to D2D3, the samples were considered positive only if the protein concentration was deemed ≥7 ng/ml*. Note that the levels of suPAR and D2D3 in D2D3 positive DN sera are similar.

(**D**) Representative images of human podocytes stained with phalloidin (F-actin) and AP5 (antibody that recognizes active form of β3 integrin). Notice that serum free media (SFM) and healthy serum (HS) do not activate β3 integrin, but that the addition of hD2D3 (2.5 ng/ml) to the HS resulted in AP5 signal. Scale bar, 5 μm.

(**E**) Scatter dot plots depicting ratio between activated β3 integrin (AP5) and overall focal adhesions determined by the paxillin staining. Notice that addition of hD2D3 to SFM activated β3 integrin on human podocytes. This demonstrates that hD2D3 activates β3 integrin on human podocytes even in the absence of any other serum proteins. For experiments with SFM, 62-79 cells were analyzed. For experiments with HS, 43-46 cells were analyzed. Data are shown as mean ± SEM. An unpaired two-tailed *t*-test was was performed to determine statistical significance (****$P < 0.0001$)

(**F**) Graph showing concentration-dependence of hD2D3's ability to activate β3 integrin on human podocytes grown in the presence of healthy human serum. 20-101 cells were analyzed per treatment. Error bar = mean ± SEM. Notice that at lower protein concentrations (1-~3 ng/ml) activation appears cooperative.

**A**

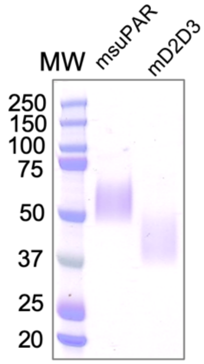


**B**

| Protein | Glyco. (kDa) | De-glycol. (kDa) |
|---|---|---|
| msuPAR-Myc | 50-70 | 36-45 |
| mD2D3-Myc **(mouse)** | 40-60 (**40-60**) | 20-30 |

**C**

| Input | BCA (ng/ml) | R&D (ng/ml) |
|---|---|---|
| mD2D3 | 1 | 0.75 |
| msuPAR | 1 | 0.56 |

**Figure S3. ELISA detects mouse suPAR and mD2D3 fragment.**
(**A**) Coomassie gel showing recombinant mouse suPAR and mD2D3 fragment that were expressed and purified from HEK-293 cells. Notice that protein purification resulted in highly pure proteins. As both proteins are glycosylated, they travel as wide bands.
(**B**) Table summarizing MWs of mouse proteins. MW of mD2D3 detected in the plasma of D2D3-Tg mice was similar to the MW of purified recombinant protein.
(**C**) Mouse-specific ELISA (R&D) detected full length suPAR and mD2D3 equally well, but with low sensitivity. BCA was used to determine protein concentrations used in ELISAs.

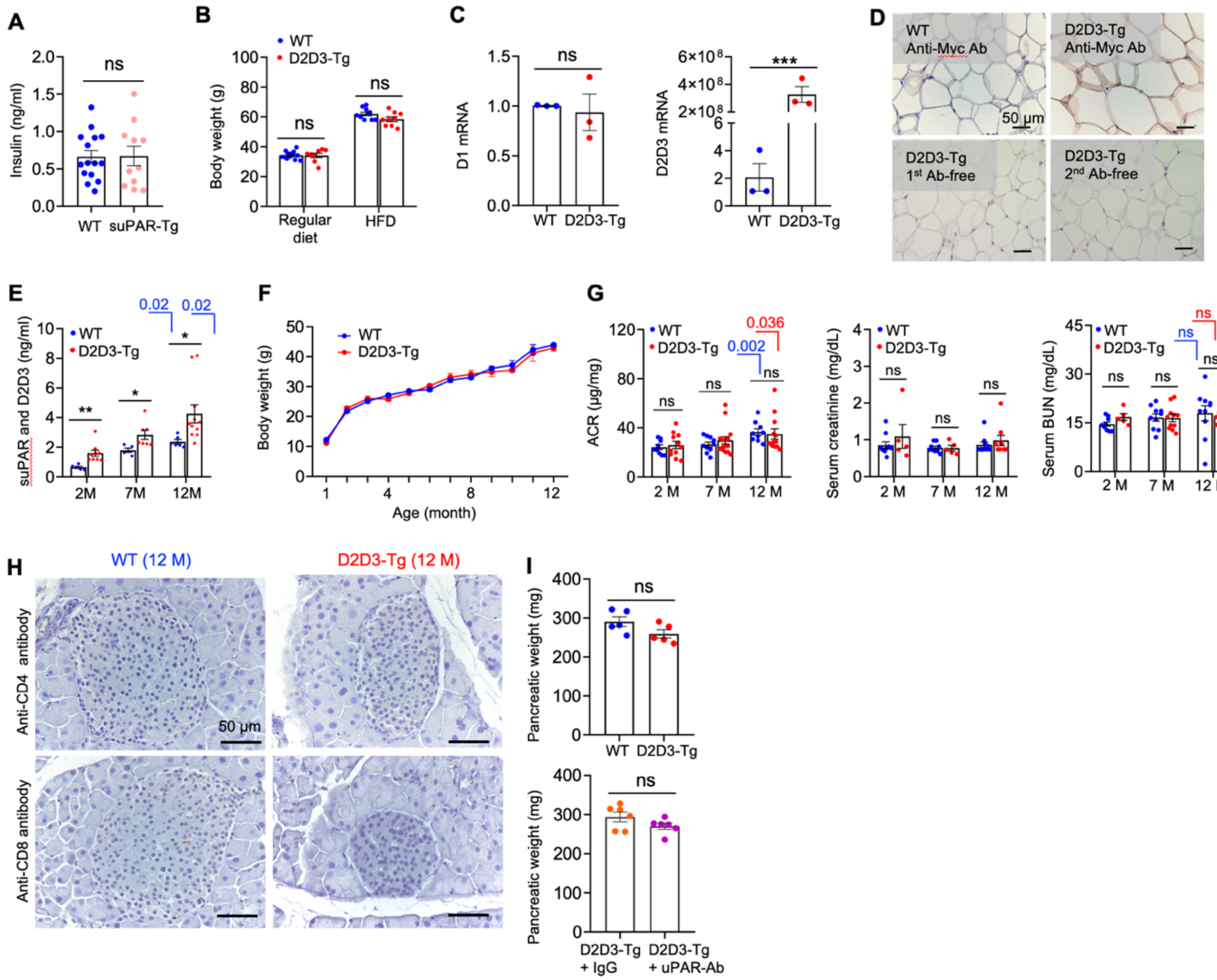

**Figure S4. D2D3-Tg mice on a regular diet do not develop glomerular injury.**

(**A**) Scatter dot plot showing fasting blood insulin levels in WT and mice expressing mouse suPAR isoform 1 (suPAR-Tg) (n = 15 for each group).

(**B**) Scatter dot plots comparing body weights of animals kept on regular diet and on high fat diet (HFD). Notice that expression of mD2D3 had no effect on the body weight of the animals regardless of their diet.

(**C**) qPCR analysis of *D2D3* and *D1* in fat tissues. *D2D3* and *D1* mRNA levels were normalized to *Gapdh* mRNA level ($n$ = 3 biologic replicates). Data show that mRNA for D2D3 was detected in the fat tissue even when animals were fed regular diet.

(**D**) mD2D3 fragment is expressed in fat tissue of animals that were fed regular diet. As D2D3-Tg carries the c-Myc tag, immunohistochemistry was performed with a rabbit anti-Myc antibody. D2D3 was observed in adipocytes of D2D3-Tg mice, but not in control mice. Tissues stained with only primary or secondary antibody were used as negative controls. Scale bar, 50 μm.

(**E**) Levels of suPAR and suPAR/D2D3 increased as animals aged. Scatter dot plots depicting protein levels determined using mouse-specific ELISA ($n$ = 5 - 11 for each group).

(**F**) Body weight of WT and D2D3-Tg mice ($n$ = 5 - 8 for each group).

(**G**) Kidney function was not affected in D2D3-Tg mice when they were fed regular diet. Kidney function was assessed by measuring ACR, serum creatinine and serum BUN levels ($n$ = 5-12 for each group).

(**H**) Immunohistochemistry of islets using anti-CD4 and anti-CD8 antibodies in 12-months old animals.

(**I**) Scatter dot plots comparing pancreatic weight between WT (n=5) and D2D3-Tg (n=5), as well as D2D3-Tg animals treated with IgG (n=6) or anti uPAR-Ab (n=6). All results were expressed as mean ± SEM. Statistical analysis was determined using 2-tailed unpaired Student's $t$ test. *$P$ < 0.05; **$P$ < 0.01; ***$P$ < 0.001. ns, not significant.

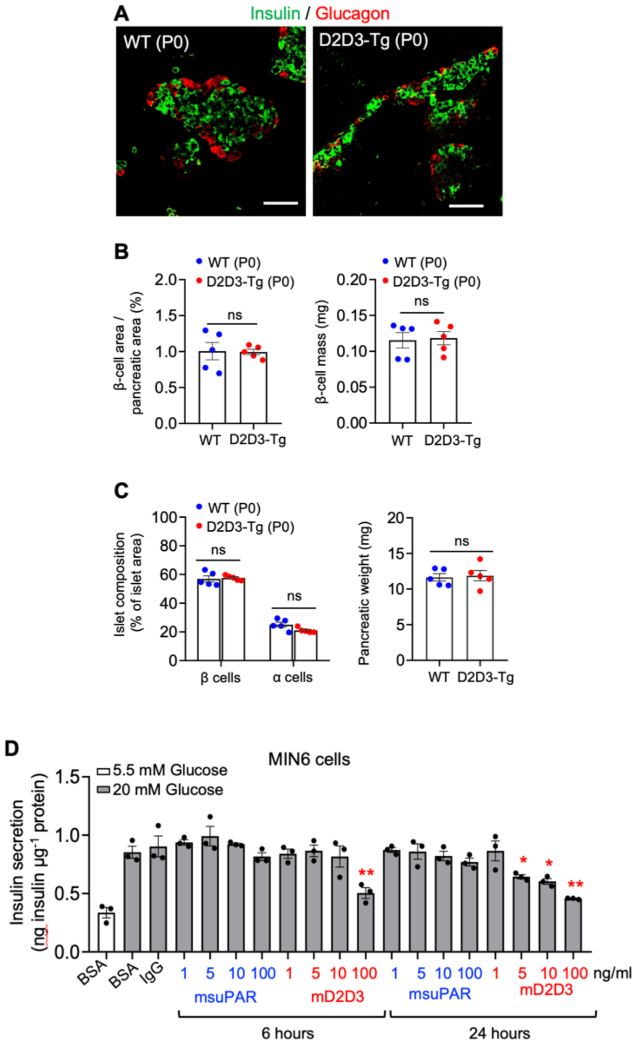


**Figure S5. Neonatal D2D3-Tg mice have normal β-cell mass.**
(**A**) Immunohistochemistry of pancreas isolated from the neonatal mice (P0) stained with anti-insulin and anti-glucagon antibodies.
(**B**) Staining shown in A was used to determine the β-cell area/pancreatic area ratio ($n$ = 5, for each group) as well as β-cell mass ($n$ = 5 for each group).
(**C**) Scatter dot plots showing Islet composition ($n$ = 5, for each group) and pancreatic weight ($n$ = 5, for each group) in neonatal mice.
(**D**) GSIS of MIN6 cells cultured in the presence or absence of BSA (100 ng/ml), IgG (100 ng/ml), mD2D3 fragment or mouse suPAR ($n$ = 3). All results are expressed as mean ± SEM. Statistical analysis was determined using 2-tailed unpaired Student's $t$ test. *$P < 0.05$; **$P < 0.01$; ***$P < 0.001$. ns, not significant.

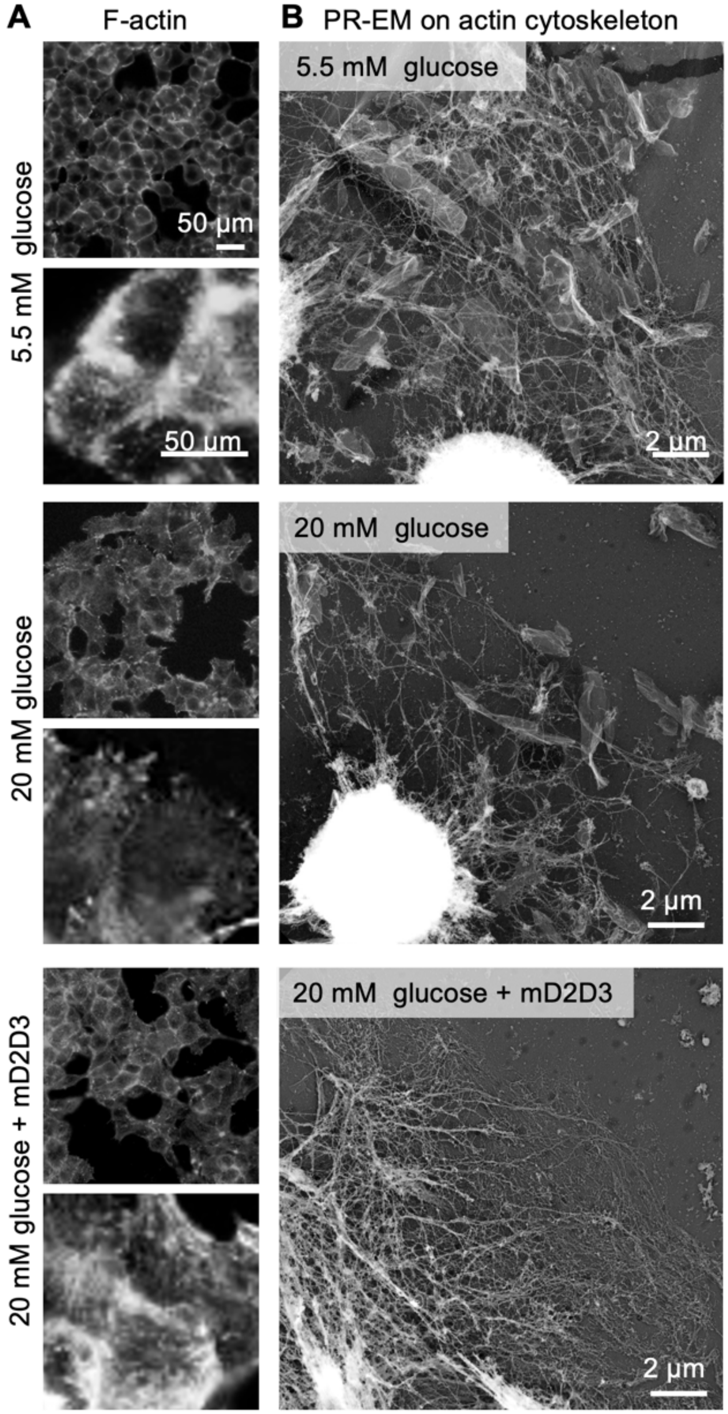


**Figure S6. mD2D3 fragment impairs glucose-stimulated reorganization of the actin cytoskeleton in MIN6 cells.**

(**A**) MIN6 cells were grown in low glucose (5mM) before being stimulated by the addition of high glucose (20 mM) in the presence or absence of mD2D3 fragment (100 ng/ml). The status of F-actin was examined by phalloidin staining.

(**B**) Representative PR-EM micrographs of MIN6 cells focusing on the organization and the status of the actin cytoskeleton. Cells were grown as described in (**A**). Notice that mD2D3 inhibited high glucose-induced disassembly of actin filaments.

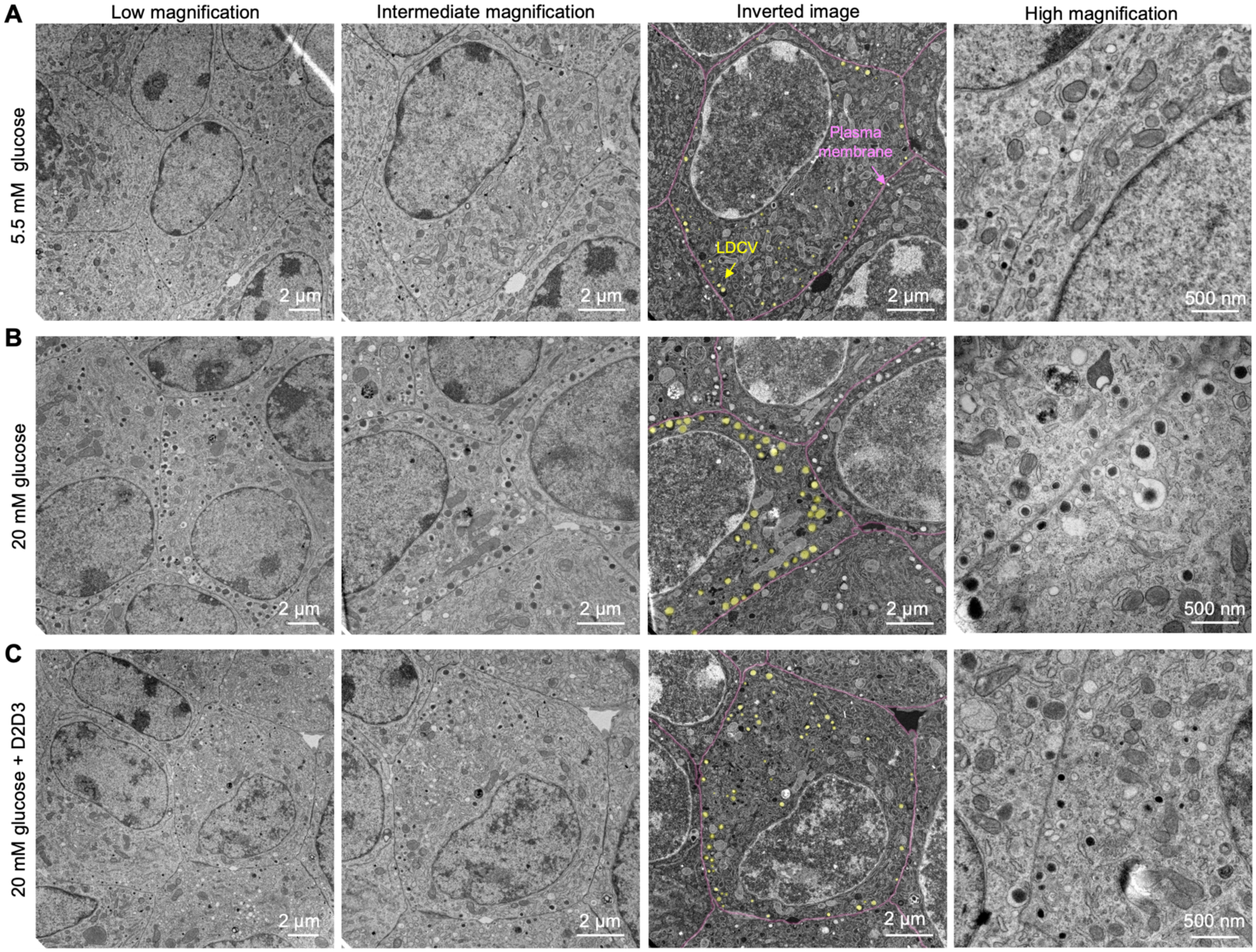


**Figure S7. mD2D3 impairs insulin granule maturation and trafficking.**
(**A**-**C**) Representative TEM micrographs of MIN6 cells. When indicated cells were grown in low glucose (5mM) before stimulating by the addition of a high glucose (5mM) in the presence or absence of mD2D3 fragment (100 ng/ml) for 24h. Images shown were captured at increasing magnifications. The plasma membrane is colored pink and large dense core vesicles (LDCVs) that transport insulin are colored yellow. Notice that LDCVs are smaller in size and less associated with the plasma membrane in MIN6 cells grown in the presence of the mD2D3 fragment.

### Materials and Methods

**Human samples:** A total of 25 human sera and matching urine samples were obtained from diabetic nephropathy (DN) patients seen at the Renal Associates, MGH by Dr. Kristin M. Corapi under MGH IRB protocol #2014P001943 in the period of 2014-2018. In addition, in the period between 2018 and 2020, we obtained 57 sera from the Mass General Brigham Biobank. Forty patients were diagnosed with DN and were not receiving insulin therapy when serum samples were collected. Forty-two patients were diagnosed with DN and were receiving insulin therapy when the serum samples were collected. Statistical analysis was performed using SPSS 27 (IBM, Armony NY). We compared the prevalence of hD2D3 (reported as a %) between patients with DN on or not on insulin using the chi-square test. To assess whether hD2D3 was independently associated with being on insulin, we used a logistic regression model with insulin therapy as the dependent categorical variable and presence of hD2D3, hemoglobin A1c % and suPAR (log-transformed base 2) as independent variables. Lastly, we computed the area under the curve (AUC) for hD2D3 and suPAR in separate models and combined to assess their ability to differentiate between those on or not on insulin therapy. We compared AUCs using the Delong test. A two-sided P-value<0.05 was adopted to indicate statistical significance.

**Reagents:** RPMI 1640 medium (11875-093), CMRL medium (21540-026); FBS (10082-147), penicillin-streptomycin (15140-122), antibiotic/antimycotic (15240-096) (penicillin, streptomycin and Amphtericin B) were from Gibco. EZ-link Micro Sulfo-NHS-Biotinylation Kit (21925), Zeba™ Spin Desalting Column, and Pierce™ IP RIPA Buffer (89901) were from Thermo Fisher Scientific. β-mercaptoethanol (M6250) and ITS (insulin-transferrin-sodium selenite) media supplement (I3146) were from Sigma-Aldrich. The reagents used for in vitro GSIS (MIN6 cells, mouse islets, and human islets) were from Sigma-Aldrich. Sodium pyruvate (Corning, 10-013-CV); protease inhibitor cocktail tablet (Roche, 11836170001); Chymotrypsin (Roche, 11 418 467 001); uPAR (R&D systems, 807-UK/CF); streptavidin Mag Sepharose beads (GE Healthcare, 28-9857-99); N-Glycanase (PROzyme, GKE-5006A).

**Cell culture:** Mouse MIN6 cells (a gift from Dr. Decheng Ren, University of Chicago) were grown as described before (Ren et al., 2014). Immortalized human podocytes were cultured according to published protocol (Saleem et al., 2002). Mouse islets were cultured in RPMI 1640 medium containing 10% FBS, 1% penicillin/streptomycin, and 50 μM β-mercaptoethanol. Human islets were cultured in CMRL medium containing 10% FBS and 1% penicillin/streptomycin.

**Antibodies:** Rabbit anti-c-Myc antibody (Sigma-Aldrich, PLA0001), guinea pig anti-insulin antibody (Abcam, ab7842), mouse anti-glucagon antibody (Sigma-Aldrich, G2654). Secondary antibody for insulin labeling was Alexa Fluor 488-conjugated goat anti-guinea pig IgG (Invitrogen) and for glucagon labeling was Alexa Fluor 594-conjugated chicken anti-rabbit IgG (Invitrogen). uPAR (R4)-BSA Free (Novusbio, NBP2-41379); Rabbit anti-uPAR (Bethyl, A304-462A); Mouse uPAR polyclonal antibody (R&D systems, AF534); anti-goat IgG HRP antibody (Thermo Fisher, HAF109); anti-rabbit IgG HRP antibody (Thermo Fisher, G-21234); AP5 antibody (Blood Center of Wisconsin); paxillin antibody Y-113 (Abcam, ab32081).

**Standard procedures:** Mouse pancreatic islets isolation was performed as described (Zhu et al., 2020). Islets were picked up manually under a dissecting microscope (Nikon Instruments Inc, Melville, NY, USA). Pancreatic β-cell area and β-cell mass were calculated as described (Zhu et al., 2020). AP5 assay using human podocytes was performed as described (Hayek et al., 2017).

**IP-coupled to western blot analysis (IP-WB):**

*Human sera and urine samples*: Human sera or urine samples were diluted (1:1) with RIPA buffer containing protease inhibitor cocktail tablet (RIPA-PI) and precleared using Streptavidin Mag Sepharose beads. Human uPAR (R4) antibody (Novusbio, NBP2-41379) was biotinylated using EZ-link Micro Sulfo-NHS-Biotinylation Kit. Biotinylated uPAR (R4) antibody was added to the precleared samples. Subsequently, Streptavidin Mag Sepharose beads were added to the rotating samples. The total immunoprecipitation (IP) time for generating samples for mass spectrometric analysis was 24 hours, and for IP-WP was 3-4 hours. The magnetic beads were washed with RIPA buffer and the bound fraction was deglycosylated using N-Glycanase. The proteins were analyzed using SDS-PAGE. Western blot analysis was performed using polyclonal rabbit anti-uPAR (Bethyl, A304-462A). Recombinant human uPAR (R&D systems, 807-UK/CF), chymotrypsin digested uPAR, or recombinant hD2D3 were used as positive controls. For mass spectrometric analysis, proteins immunoprecipitated from patient sera or urine were deglycosylyed and subjected to SDS PAGE and the protein bands were excised from the gel and submitted to the Taplin Biological Mass Spectrometry Facility at Harvard Medical School.

*Mouse sera*: Mouse sera were diluted, precleared and IPed as explained for human sera above. The antibody used to IP mouse proteins was the polyclonal mouse uPAR antibody (R&D systems, AF534). Recombinant mouse suPAR and recombinant mouse D2D3 were used as positive controls. Western blot was performed using rabbit anti-c-myc antibody (Sigma-Aldrich, PLA0001) and rabbit IgG-HRP antibody.

**β3 integrin activation assay using human podocytes:** The experiments were performed as described elsewhere (Hayek et al., 2017). Where indicated, the cells were treated with serum-free media ± hD2D3 for one set, and HS ± hD2D3 for another set. Images were acquired using a Zeiss microscope. The intensity of AP5 or paxillin was quantified using Fiji, ImageJ (NIH). Non-specific nuclear staining was disregarded while performing quantitative analysis. Data are represented as a ratio of AP5/paxillin intensity relative to the control. An unpaired two-tailed *t*-test was performed using Prism (GraphPad) to compare the treatments.

**Cloning, expression and purification of recombinant proteins:** Gene encoding mouse uPAR isoform 1 (GenBank NM_011113.4) was amplified using total RNA isolated from cultured mouse podocytes using the forward (5′-*AAGCTT* CTA CAG TCA CAT GGT CAG GGC ATC-3′) and reverse (5′-*GAATTC* TCA CAG ATC CTC TTC TGA GAT GAG-3′) primers. The PCR products were digested with the restriction enzymes *Hind*III and *Eco*RI, and subcloned into the pSecTag2A vector containing C-terminal Myc/His tag (Thermo Fisher, V90020). pSecTag2A-suPAR/D2D3 plasmids were transiently transfected into the FreeStyle™ 293-F cells (Thermo Fisher, 12347-019). Recombinant proteins were purified from the culture medium using Pierce anti-c-Myc agarose (Thermo Fisher, 20168) based on the manufacturer's protocol.

**Mice:** Mice expressing full length suPAR (suPAR-Tg) was previously described (Hahm et al., 2017). Mice expressing the D2D3 fragment (D2D3-Tg) was generated at the Transgene Facility, University of Miami. DNA encoding C-terminal D2D3 fragment of mouse suPAR isoform 1 (https://www.uniprot.org/uniprot/P35456), corresponding to NM_011113.4 in GenBank and covering amino acids 117-298 of the mature was placed under control of the aP2 promoter cassette (Wang et al., 2010) to achieve adipocyte-specific expression of the fragment. In addition, DNA encoding secretion signal peptide (Igκ) (METDTLLLWVLLLWVPGST**GD)** was placed at the N-terminal alanine 117 of D2D3 fragment to ensure that the fragment is secreted into the circulation. The signal peptide is post-translationally cleaved between glycine (G) and aspartic acid (D, shown in blod). Furthermore, DNA encoding GPI-anchor was replaced with DNA encoding a Myc-tag (see also Figure S2B).

Genotyping-positive founder mice (D2D3-Tg) were backcrossed to C57BL/6 mice for at least 5 generations to establish the colony. D2D3-Tg mice were viable and fertile. Mice were maintained on either a regular diet or high-fat diet (HFD). All animal experiments were carried out according to the NIH Guide for Care and Use of Experimental Animals and approved by the Rush University Institutional Animal Care and Use Committee (IACUC) protocol #19-014.

**Real-time RT-PCR (qPCR):** RNA was isolated with TRIzol (Thermo Fisher, 15596-026) and cDNA was generated using high-capacity cDNA reverse transcription kit (Thermo Fisher, 4368814). qPCR was performed using SsoAdvanced Universal SYBR Green Supermix (BioRad, 172-5271). All PCR data were normalized by the *Gapdh* gene expression level. qPCR primer sequences were as follows: *D2D3* (including part of the Myc tag), forward: 5′-GAGTTATACCGTAAGAGGCT-3′, reverse: 5′-CAGATCCTCTTCTGAGATGAGT-3′; *D1*, forward: 5′-AGGACCTCTGCAGGACTACC-3′, reverse: 5′-ATGGAGCCCATGCGGTAAC-3′.

**Immunohistochemical staining:** Paraffin-embedded fat or pancreas sections (5 µm thick) were deparaffinized in xylene and rehydrated using graded ethanol series (100%, 95%, 70%), followed by rinse in distilled water. Antigen retrieval was carried out by boiling the slides in a microwave in sodium citrate solution (pH 6.0). After incubation with a blocking solution (2% BSA, 0.3% Triton X-100 in PBS) for 1 hour at room temperature, the sections were stained with anti-c-Myc antibody for detection of D2D3 fragment (1:2000), anti-insulin antibody (1:500), or anti-glucagon antibody (1:300) followed by secondary antibody. Images were acquired using LSM 700 confocal microscope (Carl Zeiss).

**Cell proliferation and apoptosis:** Pancreatic sections from each group of mice were stained with antibodies against insulin or Ki67 as previously described (Zhu et al., 2020). For quantitative analysis, ImageJ (NIH) was used to count the number of DAPI positive cells within the insulin staining area. At least 1000 insulin-positive cells per mouse were counted and Ki67-positive cells were determined. Ki67-positive cells were normalized to total insulin-positive cells in the same area. Cell survival was evaluated using the In Situ Cell Death Detection Kit (Roche Applied Sciences, 11684795910) following the instructions from the manufacturer. Briefly, after dewaxation, antigen retrieval was performed using proteinase K (10 µg/ml in 10 mM Tris/HCl, pH 7.4) at room temperature for 15 min. The slides were washed twice with 1× PBS and incubated in TUNEL reaction mixture (50 µl of Enzyme solution plus 450 µl Label Solution) for 60 min at 37 °C. The pancreas sections were further stained with an antibody insulin and DAPI. The sections were examined using an LSM 700 laser scanning fluorescence confocal microscope running ZEN software (Zeiss). TUNEL-positive cells were normalized to total insulin-positive cells in the same area as what we did for Ki67 quantification.

**Assessment of renal function and insulin levels:** Urinary albumin was measured using mouse albumin ELISA (Bethyl Labs, E99-134), and creatinine was measured using an enzymatic assay kit (Cayman Chemical, 500701). Albumin-to-creatinine (ACR) ration was calculated. Kidney function was determined by measuring blood urea nitrogen level (BUN) using BioAssay Systems (DIUR-500) and serum creatinine levels were determined using Crystal Chem assay (80350). Pancreas function was determined by measuring levels of C-peptide using mouse C-peptide ELISA kit (90050) and insulin levels using ultra-sensitive mouse insulin ELISA kit (90080) from Crystal Chem. Levels of IL-6 were measured by R&D Systems (M6000B) and CRP were measured using Crystal Chem assay (80350).

***In vivo* GTT and GSIS assays:** For both assays, animals fasted overnight. Glucose was administered by intraperitoneal injection at concentration of 2 g/kg body weight. Blood samples were taken at indicated times via tail nick as the indicated time points. Levels of blood glucose and serum insulin were measured using a glucose meter (Bayer HealthCare) and ELISA assay (Crystal Chem, 90080), respectively.

***In vitro* GSIS assays**: GSIS was performed on MIN6 cells and isolated mouse or human islets. MIN6 cells were maintained in the culture medium containing 1g/L D-glucose. Indicated concentrations of recombinant mD2D3 or mouse suPAR were added to the cells for 24 hours. Before initiation of GSIS, cells were washed twice with KRBH (137 mM NaCl, 4.7 mM KCl, 1.2 mM $KH_2PO_4$, 1.2 mM $MgSO_4$, 2.5 mM $CaCl_2$-$2H_2O$, 25 mM $NaHCO_3$, 10 mM HEPES) supplemented with 0.2% BSA and 1 g/L (5.5 mM) D-glucose. Cells were placed in a low glucose KRBH media (5.5 mM glucose, 0.2% BSA) for 30 min at 37°C, followed by KRBH (0.2% BSA) media containing either 5.5 mM or 20 mM glucose for one hour at 37°C. For mouse islets, after overnight culture in 5.5 mM RPMI, size-matched islets were treated with BSA, IgG, mD2D3 or suPAR protein as indicated in the figure legends. Islets were then placed into KRBH buffer containing 2.8 mM glucose for one hour, and subjected to 2.8 mM and 16.7 mM glucose stimulation. For human islets, which were obtained from the UVA Islet Microfluidic Laboratory of the University of Virginia or ADI IsletCore Alberta Diabetes Institute of the University of Alberta (Edmonton, Canada), after overnight culture in CMRL medium, islets were incubated in 10% of human serum for 4 days.

When indicated, 10 ng/ml of human recombinant suPAR or hD2D3 fragment was added to human sera. GSIS was initiated by the end of the $4^{th}$ day as described for mice islets above. Released insulin insupernatants were determined using mouse insulin ELISA kit (Crystal Chem, 90080) or human insulin ELISA kit (Crystal Chem, 90095). The total protein was extracted by sonication for 2 minutes in 500 μl acid/ethanol (0.18 M HCl in 95% ethanol) solution as previously described (Zhu et al., 2020).

The study protocol regarding the work with human islets was approved by the Institutional Review Board of Rush University Medical Center (IRB protocol #14051401-IRB01-AM07).

**Antibody treatment:** 2-month-old D2D3-Tg mice were treated twice a week with 0.5 mg $kg^{-1}$ body weight of mouse uPAR antibody (R&D Systems, AF534), intraperitoneally injected, for four weeks.

**Extracellular flux analysis:** Experiments were performed as described in (Altintas et al., 2021). Cellular oxygen consumption rates (OCR) and extracellular acidification rates (ECAR) were detected by Seahorse XFe24 Analyzer (Agilent Technologies, Santa Clara, CA) using the Cell Mito Stress Test and Glycolysis Stress Test kits (both from Agilent), respectively. Mouse insulinoma 6 (MIN6) cells ($6\times10^4$) were seeded onto each well of a 24-well assay plate (Agilent) and allowed to attach overnight and grow for 24 h in the regular culture medium. Then, the medium was replaced with the low-glucose (2.8 mM) culture medium and cells were treated with 10 and 100 ng/ml D2D3 fragment of mouse D2D3 protein for 24 hours. On the day of the assay, the medium was replaced with the bicarbonate-free XF DMEM assay medium, pH 7.4 (Agilent) supplemented with 1 mM sodium pyruvate, 2 mM glutamine, and 2.8 mM glucose to optimize the respiration condition. The plate was incubated in a non-$CO_2$ incubator for 30 to 60 min before being transferred to the analyzer. For the mitochondrial respiration analysis, the first three measurements of OCR were recorded under basal conditions. To stimulate the cellular oxygen consumption, 175 mM glucose solution was injected (final concentration in each well: 20 mM) and OCR was measured for another 60 min. This was followed by the sequential injections of 10X concentrations oligomycin (complex V inhibitor; final concentration in each well: 1.0 μM), FCCP (uncoupler; 1.25 μM), rotenone and antimycin A (inhibitors of complex I and complex III, respectively; 0.5 μM, each). Three readings of OCR were recorded after each injection and all recordings were 8 minutes apart. This protocol allowed for an estimation of the basal respiration, proton leak, ATP production, maximal respiration, spare respiratory capacity and non-mitochondrial respiration rates as explained previously [Altintas MM et al., 2021]. For the glycolysis assay, XF DMEM assay medium was supplemented with only 2 mM glutamine whereas glucose (10 mM), oligomycin (1 μM) and 2-deoxy-glucose (2DG; 50 mM) were sequentially injected after basal ECAR readings. Three readings of ECAR were recorded after each injection and all recordings were 8 minutes apart. Basal acidification (ECAR prior to glucose injection), glycolysis (difference between ECAR before and after addition of glucose) and non-glycolytic acidification (residual ECAR after addition of oligomycin and 2DG) rates were computed to characterize glycolysis. Extracellular flux readings were normalized to protein content in each well. Therefore, OCR and ECAR measures are represented as pmol/min/μg protein and mpH/min/mg protein, respectively.

**Transmission Electron Microscopy (TEM) of kidney samples:** Mouse renal tissues were cut into 2×2 mm pieces. The tissue was fixed in Trump's fixative (EMS, 11750), dehydrated with graded ethanol, dried using a 850 Critical Point Dryer (EMS), gold-coated on a Cressington 108 Auto Sputter Coater (Ted Pella), and post fixed in 1% $OsO_4$ for 1 hour on ice. Then, the tissue was washed in 0.1 M cacodylate buffer, dehydrated, and embedded in Epon812 (EMS). Ultrathin (70 nm) sections were collected onto Formvar-coated Ni slot grids (EMS) and stained for 15 min in 5% uranyl acetate and 0.1% lead citrate. Electron micrographs were taken with Sigma HDVP Electron Microscope (Zeiss).

**Platinum Replica electron microscopy (PR-EM) and ultrathin sections electron microscopy (UTS-EM) on MIN6 cells:** MIN6 cells were grown on poly-L-lysine (Sigma, P4707) coated glass coverslips for PR-EM or ACLAR plastic sheets for UTS-EM. GSIS was performed as described above. For PR-EM, cells were detergent extracted and processed as described (Svitkina, 2016). Samples were dehydrated in a critical point dryer Autosamdri-815 (Tousimis) and coated with a 2 nm layer of platinum and stabilized with 5 nm of carbon using EM ACE600 sputter coater (Leica). The platinum-carbon replica was released from the glass coverslips on 10% hydrofluoric acid and picked up on to EM grids.

For UTS-EM, cells were detergent-extracted and fixed with 2.5% glutaraldehyde, 1.25% paraformaldehyde and 0.03% picric acid in 0.1 M sodium cacodylate buffer, pH 7.4 for 1 hour at room temperature. Then, cells were rinsed in in 0.1 M sodium cacodylate buffer three times, followed by post-fixation with 1% osmium tetroxide, $OsO_4$ and 1.5% potassium ferrocyanide, $K_3(Fe(CN)_6)$ for 1 hour at room temperature. Samples were dehydrated the same as for PR-EM, subsequently embedded in TAAB Epon (Marivac Canada Inc) and polymerized at 60°C for 48 hours. After polymerization the ACLAR was peeled off and ultrathin sections (about 50-70 nm) were cut on a Reichert ultracut S microtome (Leica), picked up on to EM grids and stained with lead citrate. Samples for PR-EM and UTS-EM were examined using a JEM 1011 model TEM (JEOL) at an acceleration voltage of 80 kV. Images were captured by CCD camera (Gatan) and presented in inverted contrast. The vesicles and membrane were colored using the brush tool with 50% opacity in Photoshop. The vesicle size was measured using the ruler tool in ImageJ software (NIH).

**Statistical analysis**: Statistical analysis was performed with Prism 5.0 software (GraphPad). Differences between 2 groups were analyzed by Student's t test. Differences between multiple groups were analyzed using two-way ANOVA. All data are presented as mean ± SEM. Difference with $P<0.05$ was considered significant.
